# A multi-omics dataset for the analysis of Frontotemporal Dementia genetic subtypes

**DOI:** 10.1101/2020.12.01.405894

**Authors:** Kevin Menden, Margherita Francescatto, Tenzin Nyima, Cornelis Blauwendraat, Ashutosh Dhingra, Melissa Castillo-Lizardo, Noémia Fernandes, Lalit Kaurani, Deborah Kronenberg-Versteeg, Burcu Atasu, Eldem Sadikoglou, Barbara Borroni, Salvador Rodriguez-Nieto, Javier Simon-Sanchez, Andre Fischer, David Wesley Craig, Manuela Neumann, Stefan Bonn, Patrizia Rizzu, Peter Heutink

## Abstract

Understanding the molecular mechanisms underlying frontotemporal dementia (FTD) is essential for the development of successful therapies. Systematic studies on human post-mortem brain tissue of patients with genetic subtypes of FTD are currently lacking. The Risk and Modyfing Factors of Frontotemporal Dementia (RiMod-FTD) consortium therefore has generated a multi-omics dataset for genetic subtypes of FTD to identify common and distinct molecular mechanisms disturbed in disease. Here, we present multi-omics datasets generated from the frontal lobe of post-mortem human brain tissue from patients with mutations in MAPT, GRN and C9orf72 and healthy controls. This data resource consists of four datasets generated with different technologies to capture the transcriptome by RNA-seq, and small RNA-seq, and supplemented this date with CAGE-seq, and methylation profiling. We show concrete examples on how to use the resulting data and confirm current knowledge about FTD and identify new processes for further investigation. This extensive multi-omics dataset holds great value to reveal new research avenues for this devastating disease.

## Background & Summary

Frontotemporal Dementia (FTD) is a devastating pre-senile dementia characterized by progressive deterioration of the frontal and anterior temporal lobes^1^. The most common symptoms include severe changes in social and personal behaviour as well as a general blunting of emotions. Clinically, genetically, and pathologically there is considerable overlap with other neurodegenerative diseases including Amyotrophic Lateral Sclerosis (ALS), Progressive Supranuclear Palsy (PSP) and Cortical Basal Degeneration (CBD)^2^. Research into FTD has made major advances over the past decades. Up to 40% of cases^3^ have a positive family history and up to 60% of familial cases can be explained by mutations in the genes Microtubule Associated Protein Tau (MAPT), Granulin (GRN) and C9orf72^4^ which has been key to the progress in our understanding of its molecular basis. Several other disease-causing genes have been identified that account for a much smaller fraction of cases^5^. Mutations in MAPT lead to accumulation of the Tau protein in neurofibrillary tangles in the brain of patients while mutations in GRN and C9orf72 lead to the accumulation of TDP-43^6^, as well as dipeptide repeat proteins (DPRs) and RNA foci in the case of C9orf72^7^.

As of today, no therapy exists that halts or slows the neurodegenerative process of FTD and to develop successful therapies there is an urgent need to determine whether a common target and therapy can be identified that can be exploited for all patients, or whether the distinct genetic, clinical, and pathological subgroups need tailored treatments. Therefore, the development of remedies relies heavily on a better understanding of the molecular and cellular pathways that drive FTD pathogenesis in all FTD subtypes.

Although our knowledge of FTD pathogenesis using molecular and cellular biology approaches has significantly advanced during recent years, a deep mechanistic understanding of the pathological pathways requires simultaneous profiling of multiple regulatory mechanisms.

Post-mortem human brain tissue is an important source for studying the disturbance of molecular processes in patients with FTD. However, systematic studies on human post-mortem brain tissue with genetic subtypes of FTD are currently lacking. Therefore, the Risk and modifying factors in Frontotemporal Dementia (RiMod-FTD) consortium has generated a multi-omics data resource with the focus on mutations in the three most common causal genes for FTD: MAPT, GRN and C9orf72.

Here, we report four datasets from the frontal lobe of post-mortem human brain from RiMod-FTD samples. We present RNA-seq, CAGE-seq, smRNA-seq, and methylation datasets from matched samples, which enable precise profiling of transcriptional dysregulations in genetic FTD subtypes. The RNA-seq dataset can be used to identify general transcriptional differences in the FTD subtypes as well as potential alternative splicing events. The regulation of the transcriptome can be studied using the other three datasets. CAGE-seq enables the detection of active or inactive promoters and thereby helps to pinpoint potentially disease-relevant transcription factors. Methylation and smRNA-seq datasets allow researchers to identify regulatory mechanisms that lead to down-regulation of certain genes. These different aspects of the transcriptome can be studied in detail due to matched samples for all these datasets.

We show that known differences and commonalities of the investigated FTD subtypes can be recapitulated with initial analyses of the datasets, such as for instance neuronal loss due to neurodegeneration. More importantly, interesting new aspects about these FTD subtypes can be easily obtained from this dataset. We therefore believe that a thorough analysis of these data from the RiMod-FTD project can reveal helpful new insights about FTD and believe that researchers will find these datasets helpful for their own studies. To make the data easily accessible, we provide a gene browser which allows to compare expression and methylation information between disease groups for specific genomic locations at www.rimod-ftd.org.

**Table 1:**
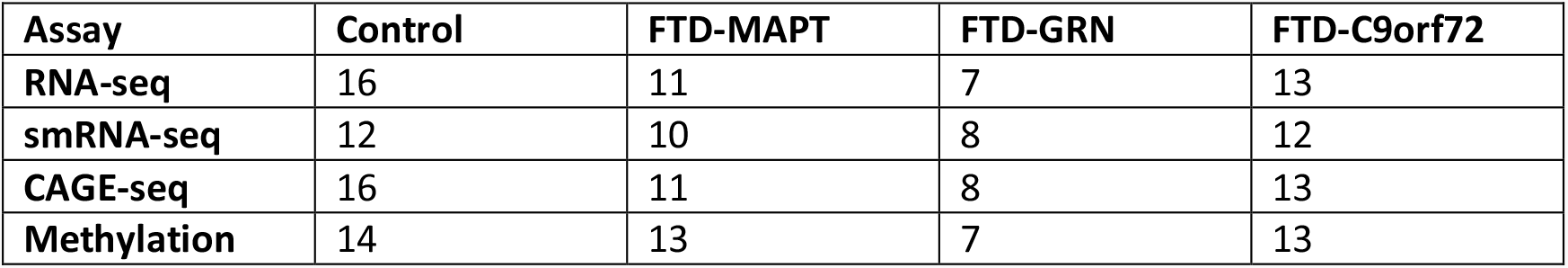
Sample numbers of the different disease groups for the available assays.

## Methods

### Donor samples employed in this study

Tissues were obtained under a Material Transfer Agreement from the Netherlands Brain Bank, and additional samples were provided by the Queen Square Brain Bank of Neurological Disorders and MRC, King College London.

GFM and GTM tissue from each subject was divided into three pieces for transcriptomic, proteomic, and epigenetic experiments in a dry-ice bath using precooled scalpels and plasticware.

### Genetic analysis

Genomic DNA was isolated from 50 mg of GFM frozen brain tissue by using the Qiamp DNA mini kit (Qiagen) following the manufacturer protocol. DNA concentration and purity were assessed by nanodrop measurement. DNA integrity was evaluated by loading 100 nanogram per sample on a 0,8% agarose gel and comparing size distribution to a size standard. Presence of C9orf hexanucleotide repeat expansion (HRF) in post-mortem brain tissues was confirmed by primed repeat PCR according to established protocols. Reported mutations for MAPT and GRN were verified by sanger sequencing.

### Transcriptomic procedures

#### RNA isolation from human brain tissue

Total RNA for CAGE-seq and RNAseq was isolated from ±100mg of frozen brain tissue with TRIzol reagent (Thermo Fischer Scientific) according to the manufacturer recommendation, followed by purification with the RNeasy mini columns (Qiagen) after DNAse treatment.

Total RNA for smallRNA-seq was isolated from frozen tissue using the TRIzol reagent (ThermoFischer Scientific). After isopropanol precipitation and 80% ethanol rinsing RNA pellet was resuspended in RNAse free water and up to 10 micrograms of RNA was incubated with 2U of Ambion DNAse I (ThermoFischer) at 37°C for 20 minutes. DNA-free RNA samples were then further purified by phenol-chloroform-isoamyl-alchol extraction followed by ethanol precipitation.

#### RNA QC

For each RNA sample, RNA concentration (A260) and purity (A260/280 and A260/230) were determined by Nanodrop measurement and RNA integrity (RIN) was assessed on a Bioanalyser 2100 system and/or Tape station 41200 (Agilent Technologies Inc.)

#### RNAseq libraries

Total RNAseq libraries were prepared from 1 microgram of total RNA from frozen brain tissue using the TruSeq Stranded Total RNA with Ribo-Zero Gold kit (Illumina) according to the protocol specifications. RNAseq libraries were sequenced on a Hiseq2500 and HISeq4000 on a 2×100 bp paired end (PE) flow cell (Illumina) at an average of 100M PE/sample.

#### CAGE-seq libraries

CAGE-seq libraries were prepared from 5 micrograms of RNA from frozen brain tissues according to a published protocol55. Libraries were sequenced on a HiSeq 2000 and/or HiSeq2500 on a 1×50 bp single read (SR) flow cell (Illumina) at an average of 20M reads/sample. On average, 21,212,923 reads were generated per sample.

#### smallRNAseq libraries

Small RNA-seq libraries were prepared from 1 microgram of total RNA from NPC-derived neurons and 300 nanograms of microglia after miRNA mimics and inhibitors transfection, using the mRNA TrueSeq Stranded kit (Illumina). mRNAseq libraries were sequenced on a NextGen550 on a 75 cycles flow cell (Illumina). Small RNAseq libraries from frozen tissue were prepared starting from 2 micrograms of total RNA using the Nextflex Small RNA-seq kit v3 (Bioo Scientific) and the NEBNext Small RNA library prep set for Illumina (New England Biolabs). Libraries were sequenced on a NextSeq550 on a 75 cycles flow cell.

#### Methylation assay

To assess the methylation status of over 850000 CpG sites in promoter, gene body and enhancer regions we have used the MethylationEPIC bead chip arrays (Illumina). Bisulfite conversion of genomic DNA, genome amplification, hybridization to the beadchips, washing, staining, and scanning procedure was performed by Atlas Biolabs (Atlas Biolabs, Berlin, Germany). Cases and controls DNAs were distributed randomly across each array.

#### RNA-seq processing and analysis

Raw FastQ files were processed using the RNA-seq pipeline from nf-core (nf-core/rnaseq v1.3)^8^, with trimming enabled. Gene quantification was subsequently done using Salmon (v0.14.1)^9^ on the trimmed FastQ files. Alignment and mapping were performed against the human genome hg38. On average, 82,165,192 reads could be uniquely mapped, which relates to on average 88.6% uniquely mapped reads per sample. In total, 59,270 transcripts could be identified. DESeq2 (v.1.26.0)^10^ was used to perform differential expression analysis. We corrected for the covariates gender and PH-value.

#### Cell type deconvolution

We performed cell type deconvolution on the RNA-seq data using Scaden^11^. For training we used the human brain training dataset used in the Scaden publication. Each ensembl model was trained for 5000 steps. Cell type deconvolution was then performed with the trained Scaden model on the RNA-seq count data. Relative changes in cell type composition were quantified by first calculating the average fractions of a cell type for all groups and then calculating the percentual change of cell fractions compared to the average control fractions. This allows to detect relative changes in cell type compositions.

#### CAGE-seq processing and analysis

Sequencing adapters and barcodes in CAGE-seq FastQ files were trimmed using Skewer (v.0.1.126)^12^. Sequencing artefacts were removed using TagDust (v1.0)^13^. Processed reads were then aligned against the human genome hg38 using STAR (v.2.4.1)^14^. On average, 16,306,077 could be uniquely mapped per sample (76% uniquely mapped on average reads per sample). CAGE detected TSS (CTSS) files were created using CAGEr (v1.10.0)^15^. With CAGEr, we removed the first G nucleotide if it was a mismatch. CTSS were clustered using the ‘distclu’ method with a maximum distance of 20 bp. For exact commands used we refer to the reader to the scripts used in this pipeline: https://github.com/dznetubingen/cageseq-pipeline-mf. In total, we could identify 47,298 different peaks.

#### smRNA-seq processing and analysis

After removing sequencing adapters, all FastQ files were uploaded to OASIS2^16^ for analysis. On average, 3,430,613 (+/- 1,365,407) reads could be uniquely mapped per sample. Subsequent differential expression analysis was performed on the counts yielded from OASIS2, using DESeq2 and correcting for gender and PH-value, as was done for the RNA-seq data. Additionally, we added a batch variable to the design matrix to correct for the two different batches of this dataset. In total, 2904 human smRNA genes could be detected.

#### Methylation data processing and analysis

The Infinium MethylationEPIC BeadChip, which consists of 866,091 CpG locations, data was analyzed using the minfi R package^17^. We removed all sites with a detection P-value above 0.01, on sex chromosomes and with single nucleotide polymorphisms (SNPs), leaving 810,290 loci for analysis. Data normalization was done using stratified quantile normalization. Sites with a standard deviation below 0.1 were considered uninformative and filtered out, leaving 170,595 sites that were used to perform a principal component analysis.

## Data Records

All datasets have been submitted to the European Phenome-Genome Archive (EGA) under the study ID EGAS00001004895.

## Technical Validation

### RNA Integrity

We only included samples with a RNA integrity value above 5. The mean RIN value is 6.9 with a standard deviation of 1.0.

### Methylation Data Quality Control and Normalization

The detection P-value has been calculated for all positions and samples. This value is calculated by comparing the total DNA signal to the background level, which is estimated using negative control positions. All samples had very low detection P-values, with a maximum value of 0.0005 (Fig. 2a). We used stratified quantile normalization to normalize the methylation signal for every position. From Fig. 2 a) and b) it is visible that this normalization can be used to even out the signal between samples in this dataset.

**Figure 1:**
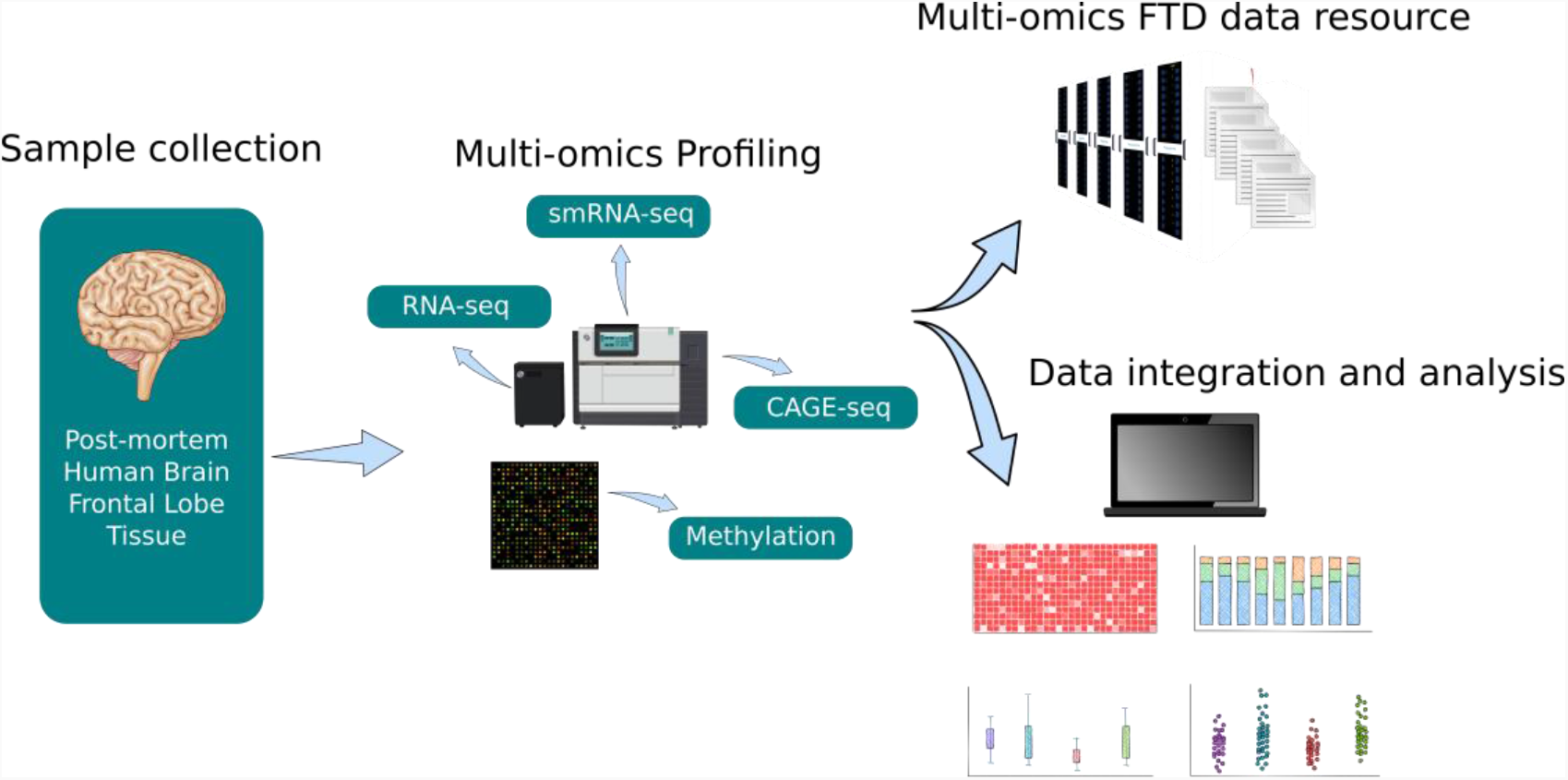
Schematic overview of the RiMod-FTD project idea. a) Samples from post-mortem human brain tissue of patients with FTD caused by mutations in MAPT, GRN and C9orf72 and from healthy controls are collected. Multi-omics profiling is performed on all samples to gain detailed insights into disease mechanisms. Integrative data analysis is performed to gain new insights and the data is made available for further studies. The goal of the RiMod-FTD project is to further extend this data resource with fitting datasets.

**Figure 2:**
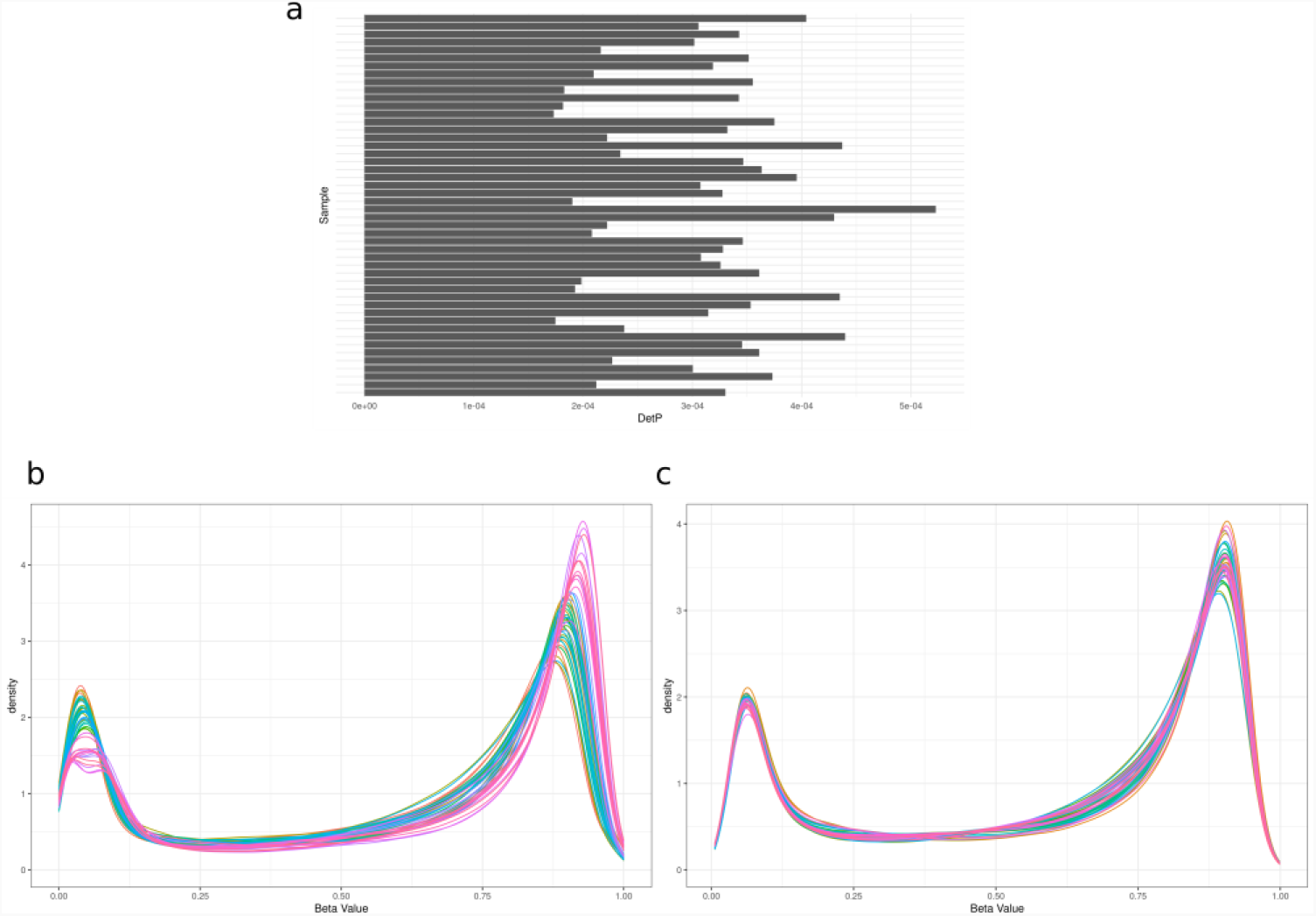
Methylation data QC statistics. **a)** Per-sample Detection P-values calculated with the minfi R-package. **b)** and **c)** Density plots of methylation beta values before and after quantile normalization, respectively.

### Principal Component Analysis of Datasets

We performed a principal component analysis (PCA) on the counts for all sequencing datasets and the M-values of the Methylation dataset (Fig. 3 a-d). As can be expected from post-mortem human brain data, the datasets are heterogeneous. However, for all assays are difference of the control samples to the different FTD subgroups can be seen. This shows that there is a clear transcriptomic difference between brains from healthy patients and from those with FTD. The PCA therefore confirms that the data is suitable for the investigation of the disease.

**Figure 3:**
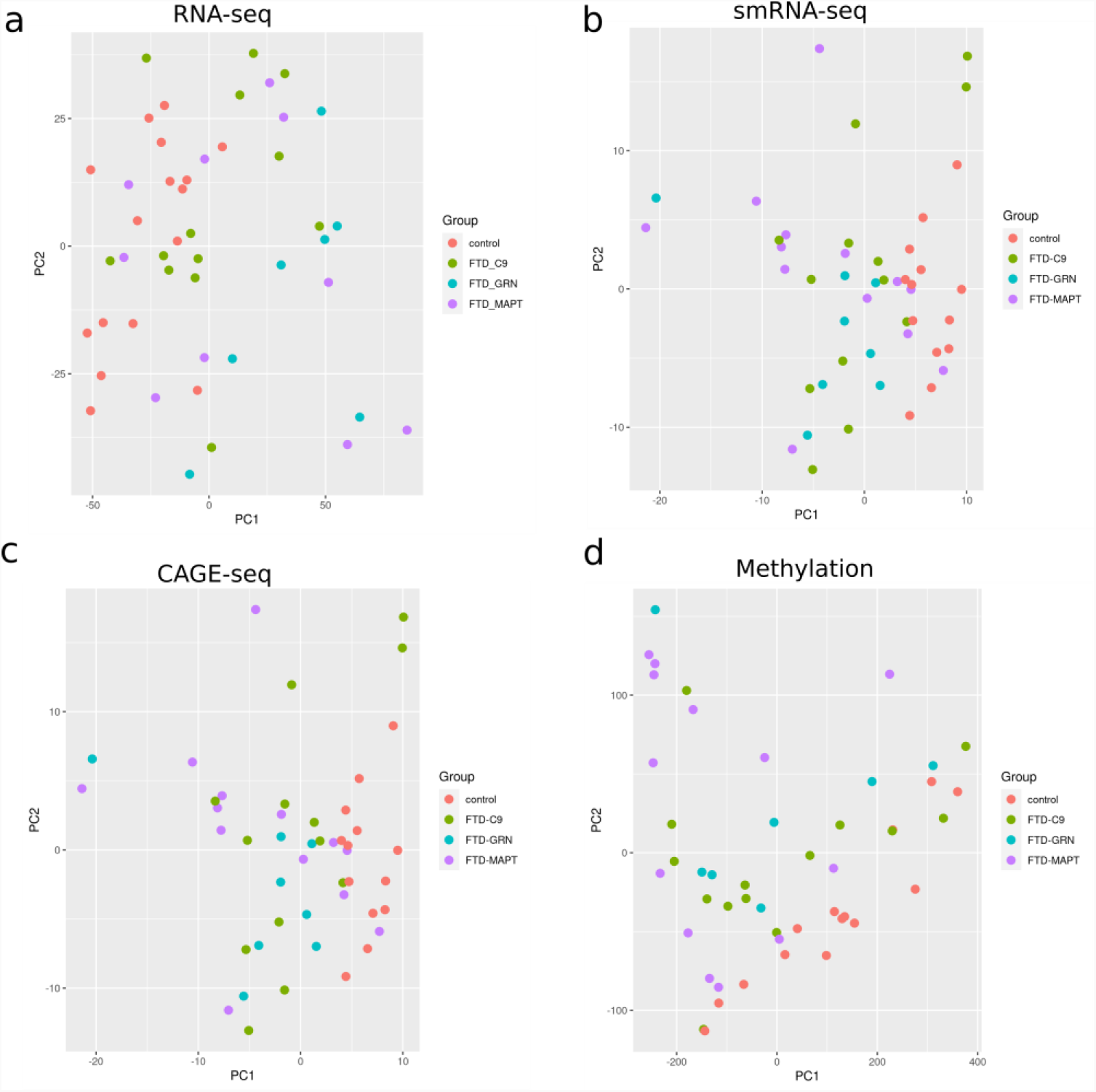
Principal component analyses of the different datasets. Shown are the first two principal components on the x and y axis. The samplesa are coloured according to the disease group. **a)** RNA-seq dataset **b)** smRNA-seq dataset **c)** CAGE-seq dataset **d)** Methylation dataset

### Cell type composition of samples

To evaluate how the predicted cell type composition of the RiMod-FTD samples reflects the disease, we performed cell type deconvolution with the RNA-seq data using the Scaden algorithm (see Methods). As expected, neuronal cells make up the largest fractions for all samples, although neuronal fractions are smaller in samples with FTD (Figure 4 b). Comparison of percentage changes of average cell type fractions compared to controls reveals that microglial fractions are particularly large in samples with FTD-GRN. Furthermore, all disease subtypes show a strong increase in endothelial cell fractions compared to controls. The decrease in neuronal cells is largely caused by a smaller number of excitatory neurons in all disease subtypes, while inhibitory neuron fractions are not decreasing. These results reflect, to a large extent, what is known about the biology of FTD.

**Figure 4:**
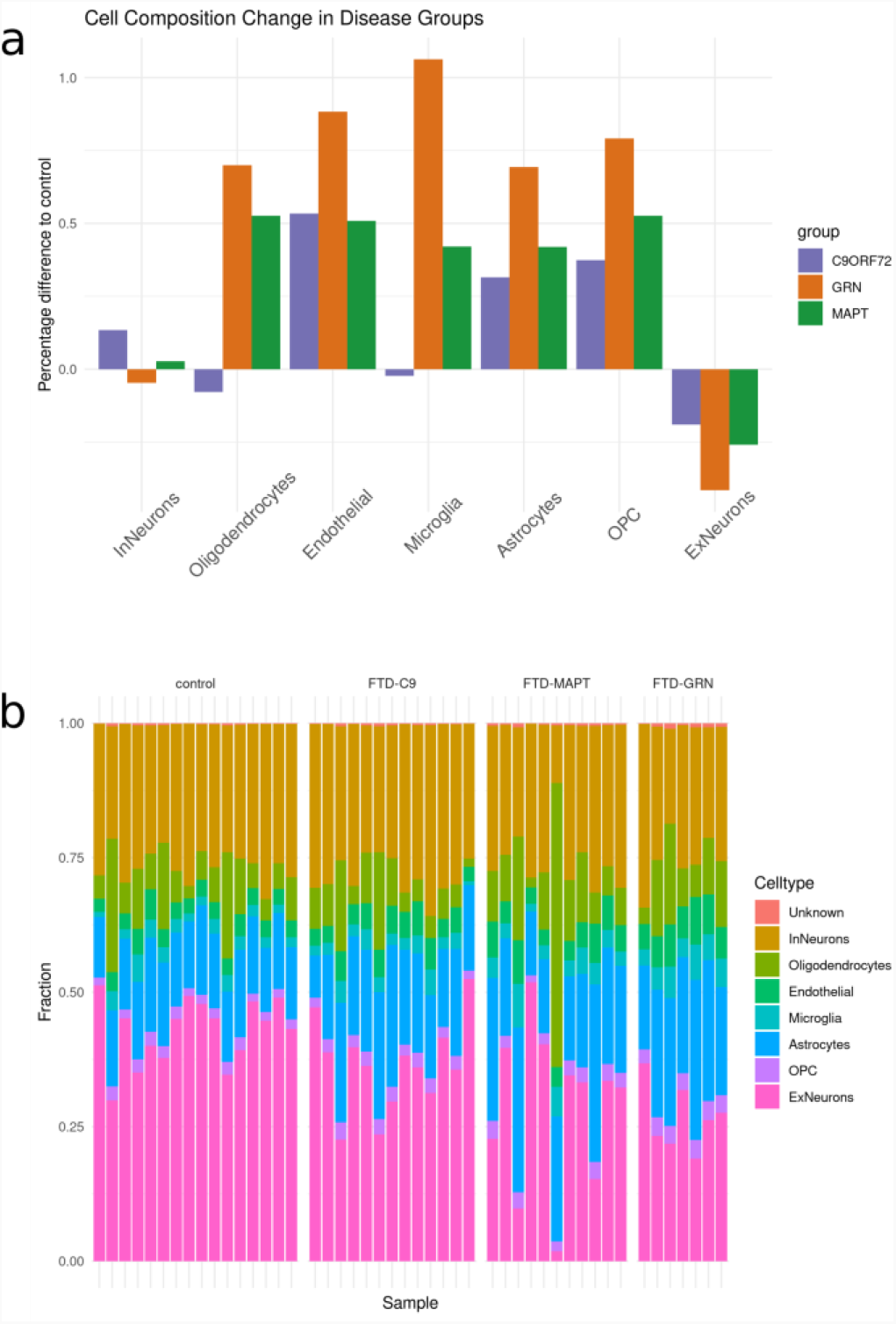
Predicted cell composition of samples and disease groups. **a)** Difference in percentage of cell compositions per group and cell type, compared to the healthy controls. **b)** Stacked bar plot showing the sample-wise predicted cell composition.

## Usage Notes

### Inspection of Assay Results in RiMod-FTD Browser

The data from the different assays of the RiMod-FTD project can be easily inspected using the RiMod-FTD Browser at https://www.rimod-ftd.org. The website allows to query the RNA-seq expression levels of single genes. It is easily visible, for instance, that GRN RNA levels are lower in brains from patients with GRN mutations, compared to the other groups (Fig. 5a). When selecting a gene, the surrounding genomic location is additionally shown with RNA-seq expression levels for genes in the region, CAGE-seq peaks and methylation levels. All data can be grouped by disease, pathology, sex, mutated gene and specific mutation (Fig. 5b-c). The browser does allow for quick inspection of FTD-related genes.

**Figure 5:**
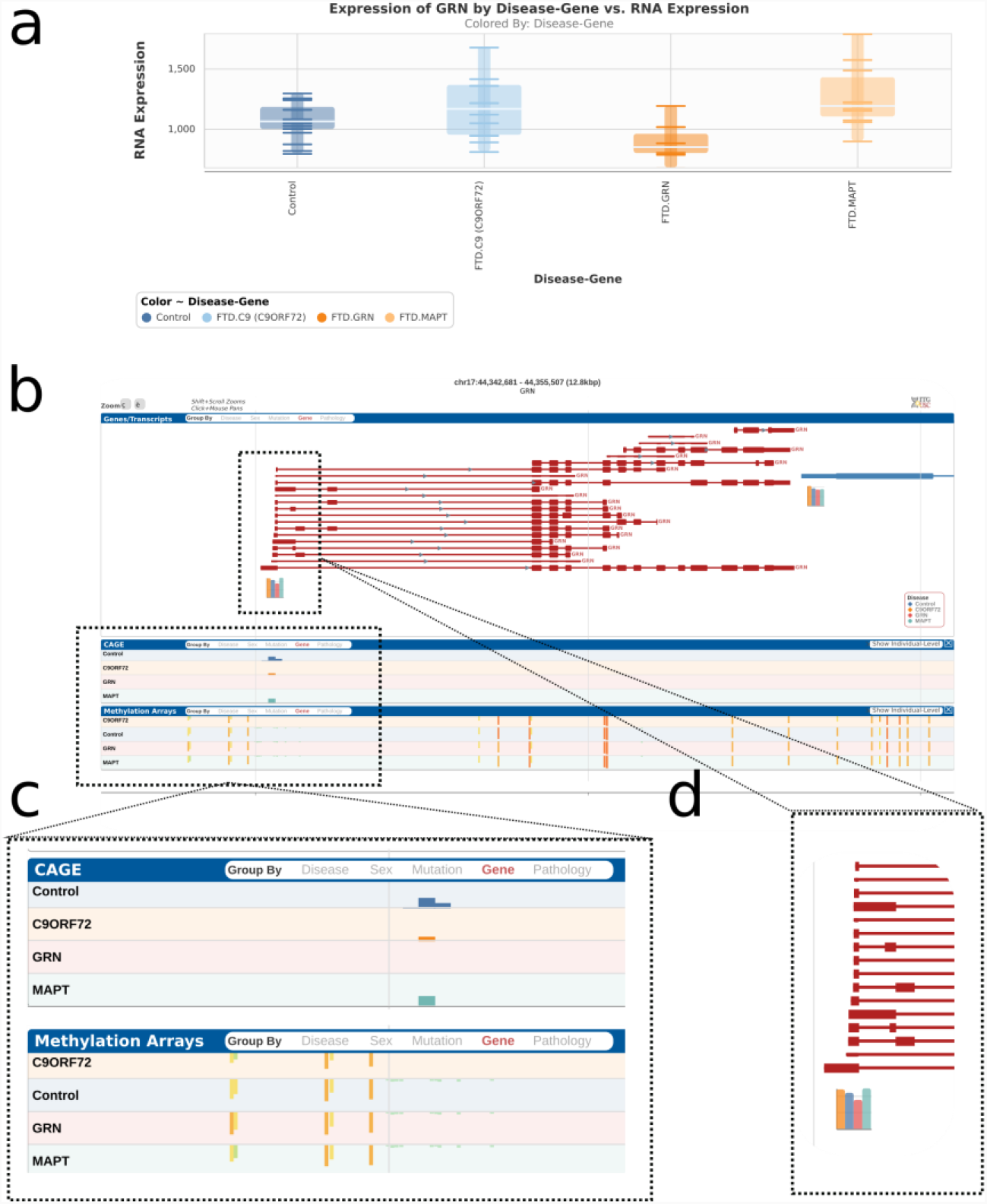
Exemplary view of RiMod-FTD genome browser. a) Normalized RNA-seq expression levels of GRN for the different disesae groups in the RiMod-FTD project. b) Browser view a selected gene (GRN). c) CAGE-seq peaks and methylation levels for the currently selected region in the browser. d) Gene expression levels for genes in the selected region of the genome browser.

## Code Availability

The code for all analyses and figures of this study has been deposited at https://github.com/dznetubingen/rimod-ftd-dataset.

## Acknowledgements

Post-mortem brain tissue was obtained from the Dutch Brain Bank, Netherlands Institute for Neuroscience, Amsterdam, and from the London Neurodegenerative Disease Brain Bank, King’s College London, London, UK. The London Neurodegenerative Disease Brain Bank is part of the Brains for Dementia Research Initiative.

## Author contributions

PH initiated and designed the project, planned, and interpreted the experiments and wrote the manuscript.

KM was involved in all bioinformatic data analyses and wrote the manuscript. AF, LK, PR and NF performed the small RNA-seq experiments.

CB, PR, NF and MC performed RNA-seq and CAGE-seq experiments. MF and TN analysed the CAGE-seq data.

BA analysed the RNA-seq data.

PR and SB planned and interpreted analyses and wrote the manuscript.

## Competing interests

The authors declare no competing interests.

## Notes

https://ega-archive.org/studies/EGAS00001004895

### Competing Interest Statement

The authors have declared no competing interest.

### Summary of Updates

The manuscript has been changed to a data descriptor format. Most data analyses will be published in a separate manuscript.

